# Lithography-less, frugal and long-term diffusion-based static gradient generating microfluidic device for high-throughput drug testing

**DOI:** 10.1101/2022.08.30.505813

**Authors:** Ketaki Bachal, Shital Yadav, Prasanna Gandhi, Abhijit Majumder

## Abstract

Drug testing is a vital step in identification of the potential efficacy of any new/existing drug and/or combinations of drugs. The conventional methods of testing the efficacy of new drugs using multi-well plates are time consuming, prone to evaporation loss and manual error. Microfluidic devices with automated generation of concentration gradient provide a promising alternative. The implementation of such microfluidic devices is still limited owing to the additional expertise and facilities required to fabricate and run these devices. Conventional microfluidic devices also need pumps, tubings, valves, and other accessories, making them bulky and nonportable. To address these problems, we have developed a method for fabricating microfluidic structures using a nonconventional technique by exploiting the Saffman-Taylor instability in lifted Hele-Shaw cell. Multi-channel structure molds with varying dimensions were fabricated by shaping ceramic polymer slurry and retaining the shape. Further using the mold thus made, polydimethyl siloxane (PDMS) devices offering static, stable, diffusion-based gradient were casted using soft lithography. We have demonstrated with COMSOL simulation, as well as using Fluorescein isothiocyanate (FITC), a fluorescent dye, that the concentration gradient can be generated in this device, which remains stable for at least 5 days. Using this multichannel device, *in vitro* drug efficacy was validated with two drugs namely-Temozolomide (TMZ) and Curcumin, one FDA approved and one under research, on glioblastoma cells (U87MG). The resulting IC_50_ values were consistent with those reported in literature. We have also demonstrated the possibility of conducting molecular assays post-drug testing in the device by microtubule staining after curcumin treatment on cervical cancer cells (HeLa). In summary, we have demonstrated a i) user-friendly, ii) portable, static drug testing platform that iii) does not require further accessories and can create iv) a stable gradient for long duration. Such a device can reduce the time, manual errors, fabrication and running expenditure, and resources to a great extent in drug testing.

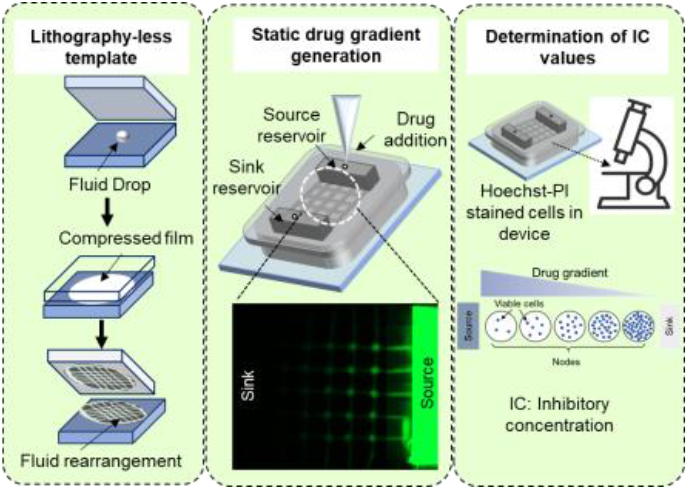

## 1. Introduction

In vitro testing is an integral part of drug discovery in which the effect of potential molecules is investigated on target cell types ^1^. Due to the advancement in genomics and molecular biology we discover numerous novel potential drug candidates almost every day which are required to be tested for their effect on various cell types ^2^.To test these drug candidates at lab-scale, drug testing assays are performed using multi-well plates in which different concentrations of the drugs are created manually. However, handling and mixing numerous different concentrations / solutions in different ratios makes it a time-consuming and labour-intensive protocol. Moreover, the working concentrations of drugs are usually in pico- to milli-molar range and hence possible manual error in dilution or error due to evaporation loss holds a serious challenge on reproducibility/reliability of the results^3,4^.To encounter these issues and fasten drug testing and drug screening procedure, high-throughput screening platforms (HTS) emerged in mid-1990s. It revolutionised drug discovery process and it is estimated that pharma companies could screen thousands of samples in a single day owing to the HTS methods. Now-a-days, due to the advancement in HTS, about 100,000 assays can be performed per day, commonly known as ultra-HTS ^4–6^.These advancements were possible due to miniaturisation and automation of conducting multi-well plate assays with the help of sophisticated robotic arms and computerised programs in a 94-well, 386-well, 1536 well or as many a s 3456- microwell plate.^5^ However, even though the HTS technology is a promising solution for sufficing the ever-increasing number drug testing and drug screening samples; most of the laboratories cannot implement them due to prohibitive cost involved.

Microfluidic devices are used for achieving high-throughput drug screening and testing. In the microfluidic approach, a precise gradient of drug concentration can be established by controlling flow/pressure conditions. In general, there are two parallel channels connected via perpendicular channels. The fluid of interest is made to flow through the parallel channels and a gradient establishes in the connecting channels via diffusion ^7^. However, gradient stability can get affected by small fluctuations in pressure or flow. Therefore, to overcome these issues for application in HTS assays, microfluidic devices were designed to create gradient by mixing and splitting fluid stream. For example, 64 cell chambers were designed for screening combination drug by *Kim et*.*al*.^8^ In this design, human bone-metastatic prostate cancer cells (PC3) were cultured and treated with drug TRAIL in combination with sensitising drug doxorubicin or mitoxantrone. The drug treatment can be either performed sequentially or simultaneously with this design. In some of HTS designs, complete channel is treated with one concentration of drug i.e., there is no drug concentration gradient. Wang et. al. reported such 576-chamber device and used it for testing cytotoxicity of five toxins on three different cell lines^9^. Though effective in practice, fabrication of such conventional microfluidic devices involves multiple fabrication steps, expensive equipments, and high level of expertise. Moreover, such microfluidic devices with fluid flow requires accessories such as syringe-pumps, tubings, customised incubators for cell-related studies etc. which make the system cumbersome and unportable. Therefore, even though multiple designs are established for microfluidic high throughput drug screening and testing, their scale-up and commercialisation is still limited and under-explored. Even though this issue can be resolved by using static devices^10^, it is difficult to maintain long-term stable drug gradient. Has C. et al., has reported one such static gradient generator which can give stable solute concentration gradients for 2 h. However, such short time period is not enough for drug response study^11^. As a result, there is no reported literature of long-term static HTS microfluidic assays.

To encounter the above drawbacks, in this work we have fabricated a static diffusion-based concentration gradient generator for long-term drug testing assay. To make the template we have exploited Saffman-Taylor instability in lifted Hele-Shaw cell (LHSC) as described by Islam T. et.al ^12^ This method has enabled us to eliminate the requirement of sophisticated lithography technique to fabricate the device, thereby making it cost-effective and frugal. Another important feature of the device is its small size, portability and stand-alone nature which has eliminated the need of bulky and expensive accessories to run the device. Using FITC we have demonstrated that the gradient generated in the device remains stable for at least 5 days. We simulated the same using COMSOL which matched well with our experimental result. Further to demonstrate the usability of this device, we estimated IC_50_ values of temozolomide (TMZ) and curcumin separately on glioblastoma cells (U87-MG). The IC_50_ values obtained from the device matched well with the same reported in the literature. In addition, we have performed immunostaining after carrying out drug-gradient study in the device to demonstrate the feasibility of post drug testing analysis.

As the operating of proposed device did not require any high level skill unlike many other flow based microfluidic devices, we envisage that this device will be an attractive drug testing platform for all categories of users (research lab, pathology clinics etc.), Use of such multi-channel platform will reduce the time, expenditure and resources to a great extent in drug testing during drug discovery, and also in devising personalised therapeutic regime.

## 2. Material and method

### 2.1. Fabrication of device

#### 2.1.1. Template preparation

In multiport LHSC we have two acrylic plates of same radii, amongst which one has holes in a particular pattern depending on the template design desire to obtain. Later, a known amount (100mg) of photo-curable viscous ceramic polymer slurry is sandwiched in between the two plates (fig 1.a-b). As the plates are separated, the holes provide preferential pathways for the air to penetrate the sandwiched ceramic polymer slurry and shape it to form the intended patterns (fig 1.c). The dimensions of square mesh can be varied by changing the initial solution volume, speed of separation of plates and distance between holes in the plate. The optimised parameters for preparation of square mesh are (i) drilled holes of 500 µm diameter, (ii) spacing between holes, (iii) squeezing fluid till 30 mm diameter and (iv) separation velocity of 5 µm/s. While the patterned ceramic polymer slurry is still wet, we can wipe unwanted part of the design to achieve desired pattern. This step makes the entire fabrication more flexible compared to the conventional techniques and renders the user with multiple number of options to edit the design at this intermediate stage. Finally, we cure the shaped ceramic polymer slurry using UV exposure (fig 1.d). Once the ceramic polymer slurry is crosslinked using UV crosslinker we use it as template to fabricate polydimethylsiloxane (PDMS)-based drug testing device (fig 1.e). Using this process, we develop template with relatively large sized square mesh network of interconnected channels of size M X N, where M= number of rows and N= number of columns. The intersections of the channels are called as nodes, which act as cell chamber in the device.

**Figure 1:**
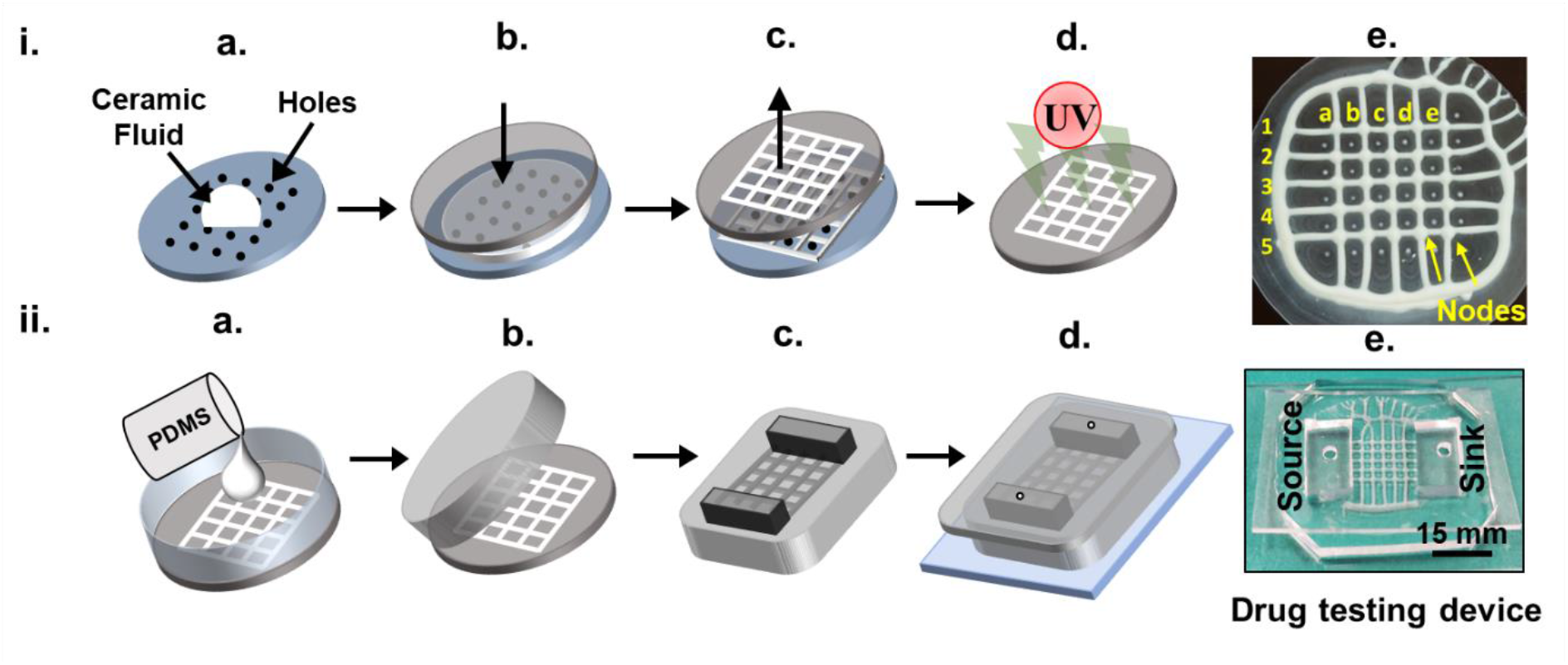
Fabrication of the device: i) Template preparation- a. Pour ceramic polymer slurry at the centre of acrylic plate with drilled holes. b. Squeeze the fluid with another acrylic plate such that all the drilled holes are covered. c. Pull the plates apart gently. d. Cure the square-mesh shaped structure thereby obtained by using UV light. e. Digital photo of fabricated template. ii) Microfluidic device preparation - a. Cast PDMS on the template and cure at 60 °C for 3 h. b. Peel the PDMS mould off the template. c. Cut the reservoirs in the mould such that the channels are connected across their length. d. Plasma bond a thin sheet (0.5 - 1.0 mm) of PDMS with two holes juxta-positioned at the middle of reservoirs on the mould and then bond this assembly on a glass slide. e. Image of the drug testing device.

**Figure 2:**
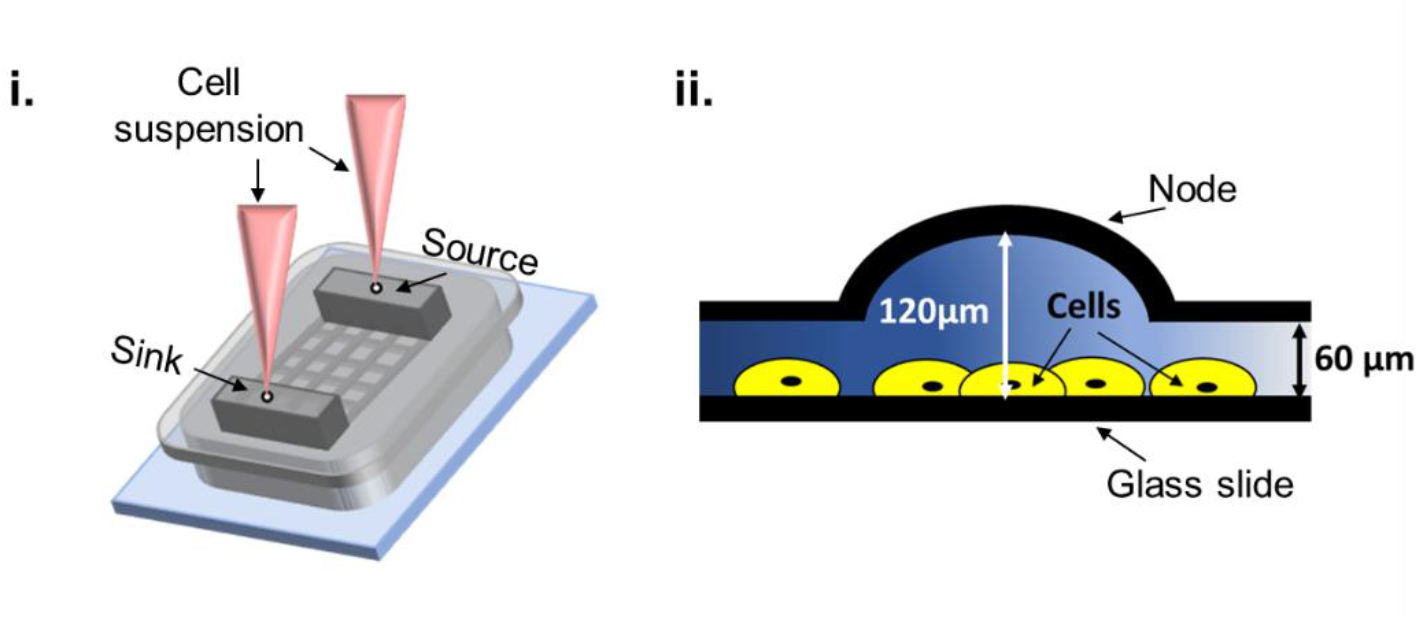
Cell seeding in the device: i) Simultaneously introducing cells in both the reservoirs. ii) schematic of cross-sectional view of the node and adjacent channel for the position of cells in the device

#### 2.1.2. Preparation of microfluidic device

Polydimethylsiloxane (PDMS) is widely used elastomer for substrate preparation as it is biocompatible, inert, nontoxic and optically clear^13^. PDMS (Sylgard 184 from Dow Corning) is mixed in various ratios ranging from 10/1 (w/w) with crosslinker and then degassed under vacuum. This uncured PDMS was then poured onto template i.e., square mesh pattern (Fig 1.ii.a) and cured at 60 °C for 12 h. The casted cured PDMS is then peeled off from template (Fig 1.ii.b). Later in this PDMS mold, reservoirs are made by cutting rectangular PDMS blocks connecting the mesh channels across their length (Fig 1.ii.c). A thin PDMS layer (with holes for inlets, juxta positioned at the centre of reservoirs) is bonded to whole device to close the reservoir in order to avoid evaporation. The PDMS mold having reservoir and a thin PDMS sheet covering reservoirs from one side is bonded on glass slide such that the channels are brought in conformal contact with glass slide for bonding (Fig 1.ii.d). At the end, we get an enclosed channel with glass base. Similarly, Figure 1.ii.e. shows the resultant microfluidic device prepared using 5 × 5 shaped template. Immediately after bonding the device we add collagen solution 25 µg/ml in the reservoirs and incubate the devices at 4 °C overnight. Post-collagen treatment the device is further used for cell seeding and drug testing.

#### 2.1.3. Template characterisation

The height of the channels and nodes were measured by using white light interferometry (WLI). WLI results in a 3D profile of the area scanned. Whereas the width of the channels and diameter of the node were calculated using the phase contrast image procured by of Evos Fl Auto (Life Technologies) fluorescence microscope (Fig 3).

**Figure 3:**
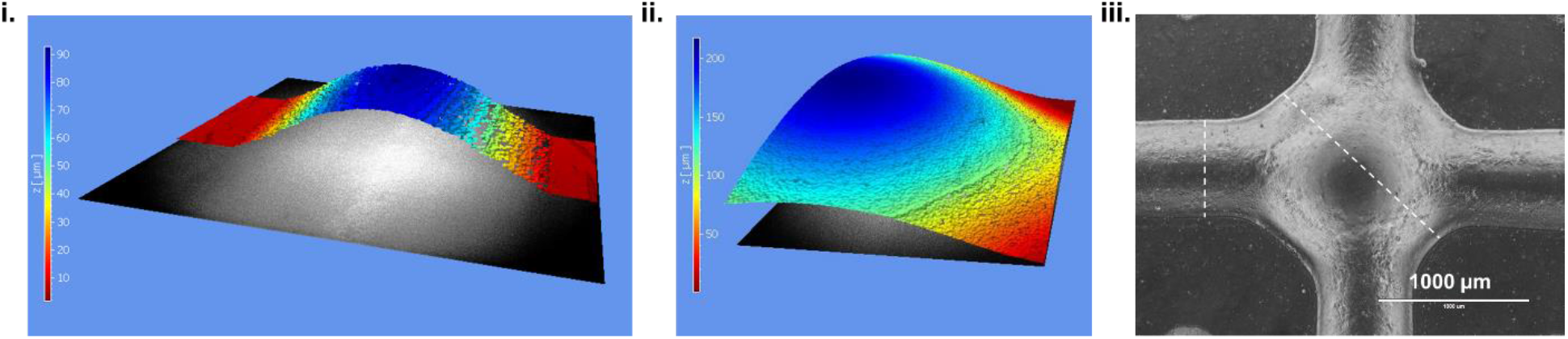
Template characterisation White light inferometer imaging for measuring- i) channel height: 60 (±30) µm ii) node height: 175 (±30) µm iii) Phase contrast image of node and adjacent channels to measure channel width: 610 (±30) µm and node diameter: 1370 (±110) µm. (represented by dashed line)

### 2.2. Gradient characterisation

#### 2.2.1. Experiment details

Fluorescein isothiocyanate (Sigma) is green fluorescent molecule, used for visualising the gradient generation and gradient stability in the fabricated microfluidic device. In the fabricated microfluidic device, the reservoirs and channels were initially filled with milli Q water. Further, we added concentrated FITC in source reservoir to achieve a 50 µM concentration in the entire reservoir. While the other reservoir with only milli Q water was referred as sink. The fluorescent intensity in the device was recorded using microscope at different time intervals (viz., 0 h, 24 h, 48 h and 72 h) and is analysed further by using ImageJ software.

#### 2.2.2. COMSOL simulation

The geometry of the device is made in COMSOL Multiphysics geometry module. The meshing is done to discretize the domains into small elements, as done in finite element method. Concentration at the outlets was simulated using COMSOL (version 5.2). Here, we used the FITC diffusion co-efficient as 4.25e-10[m^2/s]^14^. Further, the diffusion co-efficient (D) of temozolomide and curcumin were identified using the formula *D* = 0.0001778 × (molecular weight)^−.75 15^. The physics used for simulation was transport of diluted species and material was set as water medium.

### 2.3. Cell culture

The U87MG cells (brain cancer cells) and HeLa cells (cervical cancer cells) were selected as model cell line for testing drug. Cells were cultured in high glucose Dulbecco’s Modified Eagle Medium (DMEM) (Himedia-AL007A) supplemented with 1% antibiotic-antifungal solution Himedia-A002), 1% L-glutamine (Gibco) and 10% Fetal Bovine Serum (FBS) (Himedia-RM1112) in the T25 Flask till it reaches 80% confluence. For trypsinisation, the cells were washed twice with DPBS (Dulbecco’s Phosphate Buffered Saline) (Himedia- TS1006) and then 1ml trypsin-EDTA 0.05% (Himedia-TCL0033) was added for a T-25 flask having confluency of 70-80%. Trypsin and cells were incubated at 37°C for 5 mins. After incubation the trypsin was neutralised with 2 ml of complete media. To get pellet of cells it was centrifuged (1000 rpm for 5 min). After centrifugation, cells were resuspended with fresh medium and counted using haemocytometer (Invitrogen).

#### 2.3.1 Cell seeding in the device

The fabricated device (as described in section 2.1.2) with collagen treatment was exposed to UV for 20 min for sterilisation before cell seeding. Then the device was rinsed gently with DPBS three times to remove the excess/unbound collagen in the channels. Approximately 120 µl of cell suspension (desired cell density) was seeded through both the reservoirs-source and sink (60 µl each reservoir). Immediately after seeding the reservoirs with respective volume of cells suspensions the PDMS lid of reservoirs was gently pressed (one reservoir at a time) for 8-10 times without introducing air bubbles inside the channels. After ensuring that all the nodes have approximately equal number of cells using microscope, excess cell suspension in the reservoirs was removed using syringe needle of gauge 2 mm and volume 2 ml. Then the device was kept in CO_2_ incubator at 37 °C for 30 mins. Once the cells were adhered to the bottom surface of channels, then the device was carefully filled with 10% FBS media, making sure that air is not trapped inside the channels.

#### 2.3.2. Drug gradient generation in the device

a. Temozolomide gradient- After cell seeding, the devices were incubated for at least 6 h at 37 °C to ensure the attachment of all the cells in the device. Then the media in test and control device (0 µM TMZ throughout the device) was gently replenished with fresh media. Whereas in DMSO control and positive control device (device with 500 µM TMZ throughout the device), the media was replaced by the media with DMSO and 500 µM TMZ (Combi-Blocks, Cat. No. OR-2567) respectively. In test device 0.25 µL from 100 mM TMZ stock solution was added to achieve 500 µM in the source reservoir (calculations based on the volume of source reservoir). The opening on the source and sink were then covered with a thin (0.2 mm) sheet of PDMS. b. Curcumin gradient- Here, we used two devices- i) test device with source reservoir having 50 µM curcumin concentration whereas sink has 0 µM curcumin concentration ii) DMSO control device. These devices were seeded with 1.0 × 10^5^ as mentioned in section 2.3.1. Later, after cells were attached in the device we added 0.3 µL of 100 µM curcumin (Sigma Aldrich, Cat. No. C1386) in source to achieve 50 µM concentration in source (calculations based on the volume of source reservoir). The inlets and outlets of source and sink where covered with thin PDMS sheet of 0.2 mm thickness. For next 48 h we allowed the curcumin gradient to form through the channels and act on the cells.

#### 2.3.3 Live-dead assay in the device

Hoechst (4′,6-diamidino-2-phenylindole) and PI (propyl iodide) were used for staining the cells in test and control devices. After 72 h of TMZ treatment, the media in the test devices and control devices was gently removed using syringe by tilting the device such that the source end is at the bottom. Later fresh media with Hoechst 33342 (Life Technologies, Cat. No. H3570), (1:20000) and PI (Sigma Aldrich- Cat. No. 81845) (1:500) was introduced in the device. After 20 min of incubation at 37°C in dark condition, fresh media was introduced in the device and imaging of staining was performed. The removal of media through the device during staining also aided the washing off of dead cells and debris remnant cells in the device. Images were taken for all 25 nodes of test and control device in Hoechst and RFP channel of Evos Fl Auto (Life Technologies) fluorescence microscope. Later these images were analysed for the total number of live cells - counted in ImageJ software. Measurements performed are:

Number of live cells = Number of Hoechst stained nuclei - Number of PI stained nuclei

%survival= (Number of live cells/ Number of cells in DMSO control) *100.

%inhibition= (100-%survival)

This percent percentage inhibition at 5 nodes was further plotted against the concentrations of drug (TMZ/Curcumin) at the respective nodes determined using COMSOL simulation (Fig 4.iv). As the concentration of the drugs changes over time at least for the initial time points (fig 6 iii and fig S4 iii), we used time averaged integrated concentration at each node in these plots. From these plots the IC_50_ value was determined using a template in GraphPad prism- agonist vs. normalised response.

**Figure 4:**
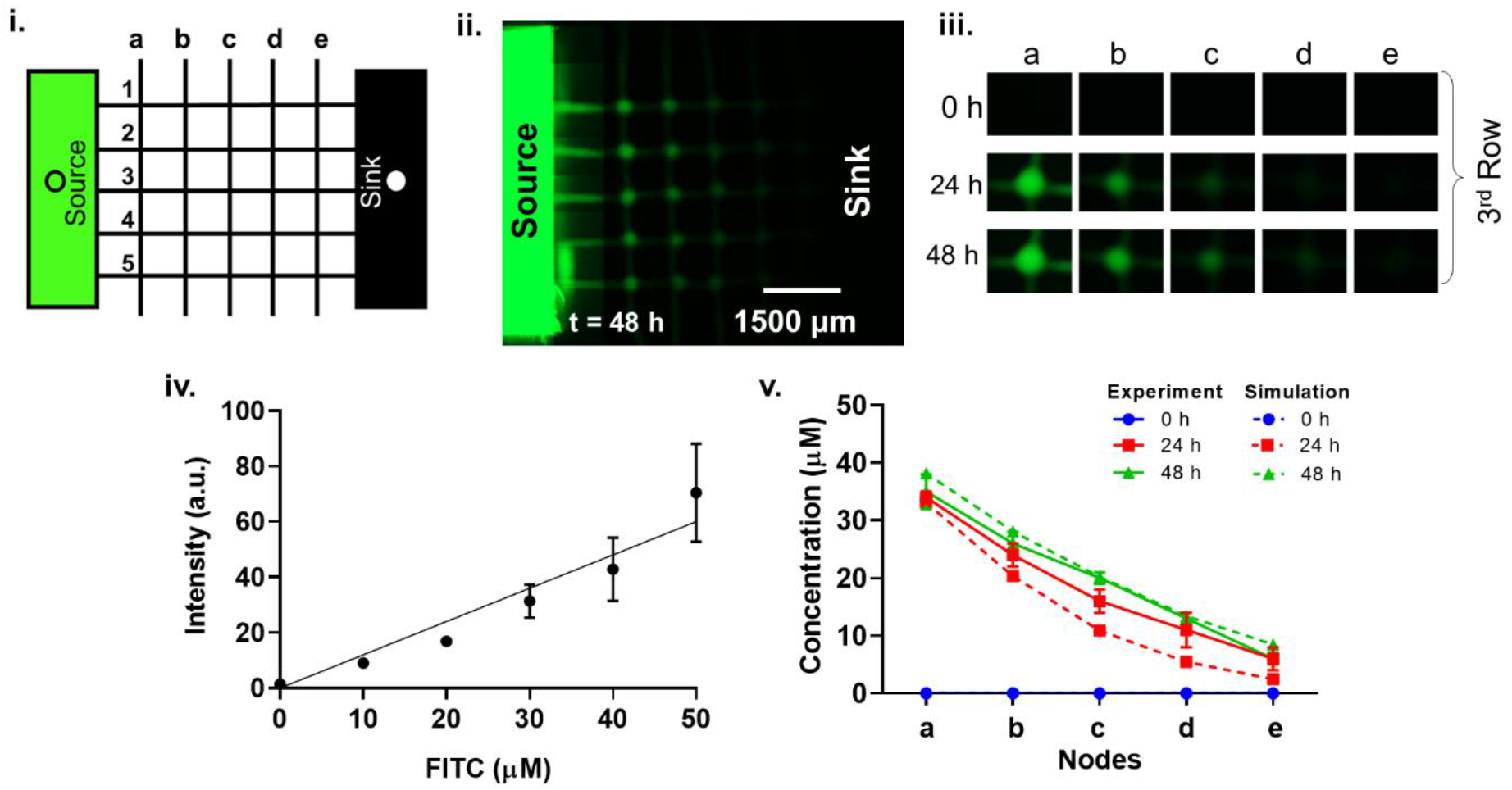
Stable gradient generation in device: i) Nomenclature used for device labelling: channels parallel to the reservoirs are labelled as a, b, c, d and e from source to sink. Channels perpendicular to the reservoirs are labelled as 1, 2, 3, 4, and 5. ii) Gradient of Fluorescein isothiocyanate (FITC) in device at 48^th^ h. iii) FITC at 5 nodes of 3^rd^ row in the device (from source to sink) at 0 h, 24 h and 48 h of the gradient generation. iv) Standard curve for FITC intensity at known concentration v) Experimental and simulation gradient profiles of FITC in device at 0 h, 24 h and 48 h.

**Figure 5:**
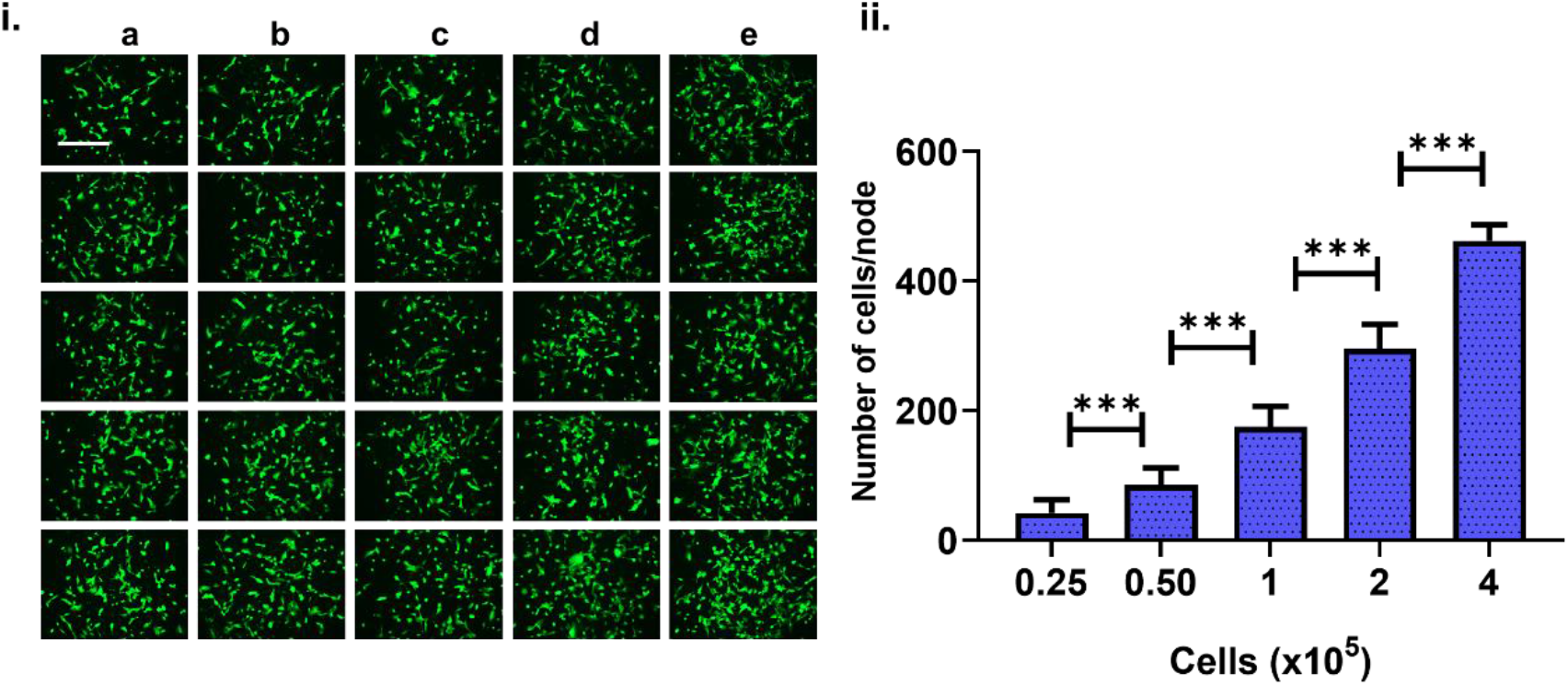
Equal distribution of cells at nodes of device: i) Calcein stained cells showing near equal distribution of live cells at 25 nodes of device with seeding density of 1.0 × 10^5^ cells. (Scale bar 400 µm) iii) Number of cells per node in the device at varying seeding densities in the device.

**Figure 6:**
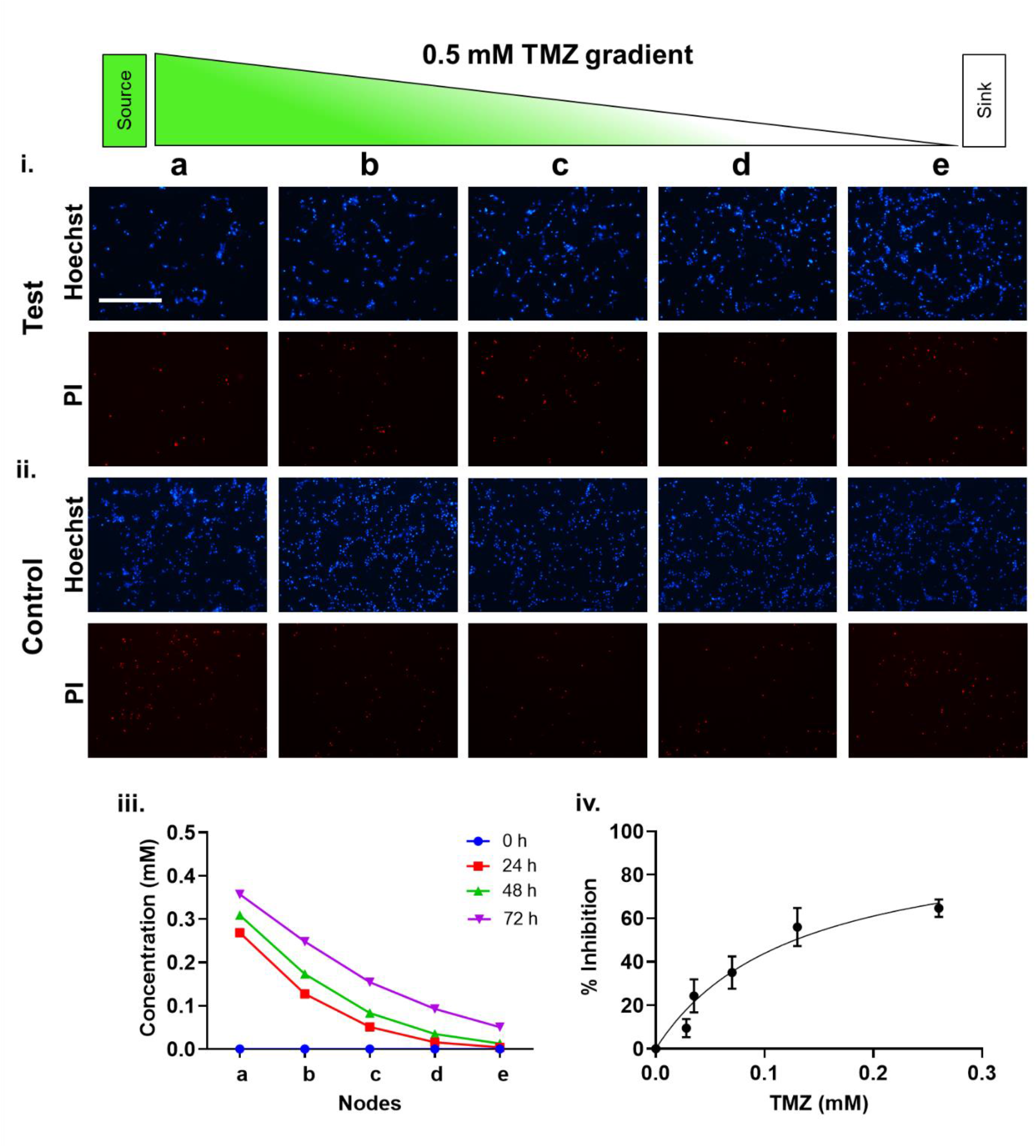
Estimation of Temozolomide (TMZ) efficacy using the device: i) Test device: Effect of TMZ gradient on U87-MG cells was determined using Hoechst and PI staining from source to sink (a-e) at 3^rd^ row in device. ii) Vehicle control device: Hoechst and PI staining at 5 nodes (a-e) of 3^rd^ row in the device. iii) Simulation graph of TMZ gradient using COMSOL at 0 h, 24 h, 48 h iv) Percentage inhibition of U87 MG cells at different concentration of TMZ at 5 different nodes from source to sink (a-e) gave IC_50_ as 0.128 ± 0.038 mM (Scale bar 400 µm).

### 2.4. Immunostaining in device

To demonstrate a post drug testing assay in device, we have performed immunostaining of microtubulin in cervical cancer cells (HeLa cells) treated with curcumin gradient. After drug gradient incubation we terminated the incubation by discarding the media and passing 0.1 % triton X-100 in DPBS for 15 sec. Immediately after this step we fixed the cells in device by flushing 4% PFA (paraformaldehyde solution in cytoskeleton buffer) through the channels for 10 min followed by washings with CSB (cytoskeleton buffer) twice and one wash with 50 mM NH_4_Cl to quench PFA. Then, we peeled off the entire PDMS structure of device, leaving fixed cells attached to the glass slide. Later cells were permeabilised with 0.5 % triton X-100 made in CSB buffer for 5 min followed by 2 washes of CSB. These cells were then subjected to blocking reagent (3 % BSA + 0.1 % triton X-100 in CSB) for 30 min at room temperature. Cells were then incubated with primary antibody against β-microtubulin raised in mouse (1:200 in blocking reagent) overnight at 4 °C. Next, we did secondary antibody staining using goat anti mouse Alexa Flour 488 (1:500) along with Hoechst 33342 (Cat. No. H3570) staining (1:10000) for 30 min at RT followed by 3 washes of CSB. All the images were captured using laser confocal imaging (Carl Zeiss) at 40X (oil immersion) magnification.

## 3. Result and discussion

### 3.1. Template characterisation

To get the dimension of the channels and nodes (i.e., intersection of branches as shown in the figure 1i) images of the template were taken using white light interferometer which gave us the height of the structures (Fig 3i and ii). The width of the channels and the nodes were found from phase contrast images taken using a light microscope. The template used in this study has the average channel height 60 ± 30 µm and width 610 ± 30 µm, whereas the nodes have average height 175 ± 30 µm and diameter 1370 ± 30 µm (Fig 3).

### 3.2. Gradient characterisation

To estimate the gradient stability, we added 0.28 ul of FITC (Fluorescein isothiocyanate) of molecular weight- 376 Da in the source reservoir of volume 600 µl resulting into a final reservoir concentration of 50 µM, as shown in the schematic of figure 4.i. The device was kept undisturbed other than during imaging. The gradient was generated due to the diffusion of FITC molecules through the static liquid present in device. Figure 4. ii. shows the stitched fluorescence image of the device showing the gradient of FITC at 48 h. Representative images of the 3^rd^ row of fabricated device at 0 h, 24 h and 48 h (from source to sink) are shown in figure 4.iii to depict the evolution of the concentration gradient inside the device. The concentration gradient was estimated by measuring the mean intensity of FITC in each node at each time point using ImageJ software and then calibrating against a standard curve (Fig. iv.). The result is shown in figure 4. v., (experimental condition represented by line graph) depicting stability of the gradient from 24 to 48 hrs. The system was also modelled in COMSOL using the transport of dilute species model which matched well with our experimental findings (Fig 4. v., simulation represented by dashed-line graph). For the simulation, we used the diffusivity of FITC in media as 4.25 ± 0.01 × 10^−6^ cm^2^ s^−1^ at 25 °C ^16^. Our results demonstrate that the device can produce multiple concentrations at different nodes, and the concentration gradient remains stable for 6 days (144 h) (supplementary figure 3), which implies the possibility of long-term experiments in the fabricated microfluidic device. We have also demonstrated that our COMSOL model can be used to predict the concentration at different nodes if the diffusivity of the diffusing molecules is known. We will use this understanding for drug testing as described in the following sections.

### 3.3. Equal distribution of cells at the nodes of device

To simplify the calculations during the drug testing assay, it is important to have nearly equal number of cells in each node of our device. Therefore, to achieve the equal distribution of cells in device, we followed the method described in section 2.3.1 to seed U87 MG cells. We seeded the cells at varying densities in the device viz., 0.25 × 10^5^, 0.5 × 10^5^, 1.0 × 10^5^, 2.0 × 10^5^, 4.0 × 10^5^ cells to understand the relationship between seeding density vs cell density at the nodes. In each node, due to its unique dome shape, cross-section increases suddenly causing a abrupt drop in the flow velocity. As a result, cells get deposited at the nodes which are then analysed later for testing cytotoxicity of the drug. These cells were stained with Hoechst and PI after 24 h of seeding in the device (supplementary figure 1). A representative image of calcein stained U87-MG cells equally distributed in the 25 nodes of device is shown in figure 5.i at 1.0 × 10^5^ cell seeding density. As the seeding density of cell/ml was increased, number of cells per node also increased as expected (Fig 5.ii, supplementary figure 1). The number of cells/nodes was quantified using Hoechst/PI images and ImageJ software (fig 5.ii). This figure gives us two informations. First, depending on the experiment, we may decide the desired number of cells/node and we may start with the required cell density in the seeding medium. Second, the error bar in the graphs show the variation in cell numbers. We observe that the variations are more prominent when a smaller number of cells/nodes are seeded. Hence, for our next experiments of drug testing, we used seeding concentration as 1.0 × 10^5^ cells which gives us a cell density/node as 175 cells with ±32 variation in cell number. To note that although 0.1 million cells are added in the device, only ∼4000 (25×175) cells are deposited in the nodes. Rest of the cells remain in the reservoir and are collected back soon after the seeding. Such process does not disturb the cells in the nodes and hence the collected cells can be used for other purposes.

### 3.5. Determination of drug efficacy using the device

Glioblastoma Multiforme (GBM) is a highly malignant and aggressive form of brain cancer, Even after the advances in medical sciences over the years, there has not been much improvement in the prognosis of the disease^17,18^. Temozolomide, a chemotherapeutic drug, is widely used for the treatment of glioblastoma ^19^. U87-MG is a glioblastoma cell line derived from malignant glioma and commonly used as the *in vitro* model cell line for GBM.^20^ As both the cells i.e. U87-MG and drug (TMZ) are well-established in the field, we used the combination to validate our device. We have used our device to determine the IC_50_ for TMZ on U87-MG. For that, U87-MG cells were seeded at a seeding concentration of 1.0 × 10^5^/ml of media in the 6 devices such that the cells get equally distributed at the nodes (method section 2.3.1). Three devices were labelled as test and other devices were used as vehicle control (with DMSO quantity equivalent to its volume to volume ratio used to make 0.5 mM TMZ), 0.5 mM control (positive control, with 0.5 mM of TMZ in the device) and 0 mM control (negative control, with no DMSO and TMZ). After cell seeding we proceeded with the protocol of drug testing assay as mentioned in section 2.3.2. After subjecting the cells to the gradient generated in the device for 72 h we terminated the experiment and performed live dead assay as mentioned in the section 2.3.3.

We observed that cell numbers, indicated by Hoechst stained- blue colour nuclei, decreased with increase in TMZ concentration as we move from sink to the source reservoir, as shown in fig 6.i and in the supplementary fig 2.i and 2.ii). Since TMZ is an anti-proliferative drug and not a cytotoxic drug, no change in the number of dead cells (PI stained cells) was observed in the test devices. Least number of live cells were observed in the positive control device (supplementary fig 2.iii, 2. iv, 2. v and 2.vi).

We further quantified the number of cells at the node of the devices. To identify the IC_50_, we simulated gradient profile of 0.5 mM TMZ using COMSOL (Fig 6.iii). Using this simulated profile, the concentrations of TMZ at five nodes (a-e) of the device at 72 h were estimated. Further the time-averaged integrated concentration at five nodes by 72^nd^ hour was plotted against the percentage inhibition at the respective nodes as described in section 2.3.3. The IC_50_ of TMZ using device was found to be 0.128 ± 0.038 mM (Fig 6.iv) which matched well with the literature ^21–25^. Further, we tested the effect of curcumin on U87-MG. Curcumin is being investigated for its anti-cancer against glioblastoma^26^. IC_50_ of curcumin derived from device is 16.6 µM ± 5.5 µM (supplementary fig 4) similar to that reported in literature ^27^. Hereby, we successfully validated our device by comparing IC_50_ of drugs found using our device with the same found using conventional drug testing methods.

### 3.5. Post drug-testing analysis in the device- Immunostaining of microtubule

Curcumin is widely known for its anti-cancerous properties and is being investigated for its mechanism of action in various types of cancers ^28^. It is shown that curcumin binds to the tubulin subunit of microtubule and disrupts it in the cervical cancer cells, thereby inhibiting cell growth ^29^. To test if the device can be used for post-drug testing molecular assays, we seeded cervical cancer Hela cells in our device at 175 cells/node seeding density. The cells were immunostained for tubulin after 48 h treatment with curcumin gradient (method section 2.3.2.b.). After the treatment, we fixed the cells in device and then peeled off the PDMS part of the device from the glass surface. This gave us fixed cells on glass slide, which were further stained using the immunostaining method described in section 2.4. As can be seen from the microscopy images, as the concentration of curcumin increases from sink to source, the distortions in microtubules also increases with maximum distorted microtubules at node “a” and least distorted microtubules at node “e”. (Fig 7). Such distortions were not observed in DMSO control device confirming that the device conditions or the vehicle did not interfere with the microtubule organisation (supplementary fig 3). With this we demonstrated that molecular assays post-drug treatment is possible in the device which can help in understanding the working mechanism of the drug and hence expanding the usability of our device.

**Figure 7:**
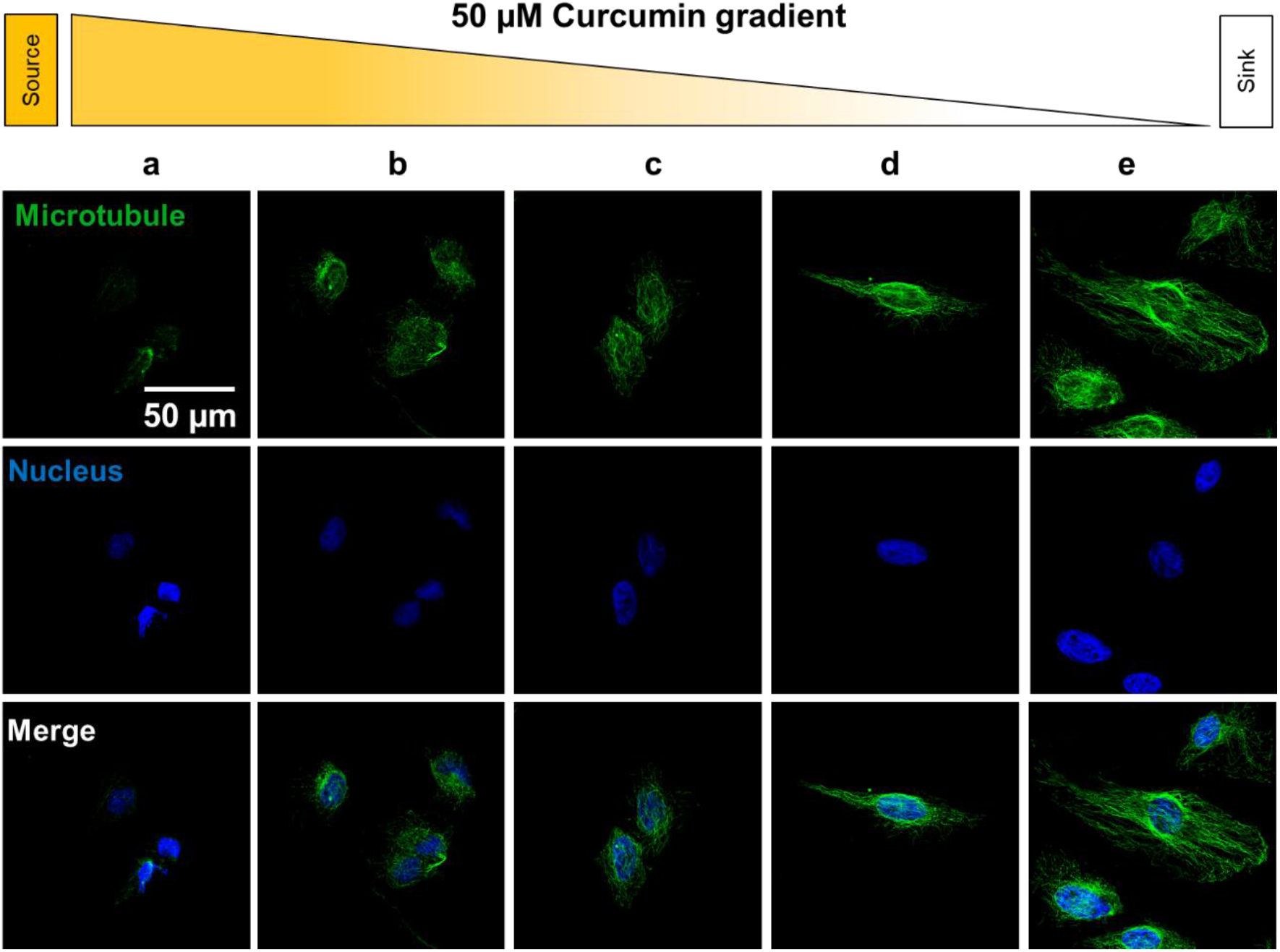
Gradient in distortions of microtubule in cells due to the effect curcumin gradient in device was captured using microtubule immunostaining in cells of test device (HeLa cells) at 5 nodes of 3^rd^ row from source to sink (a-e) reservoir. Microtubule (green), Nucleus (blue).

### 3.6. Scalability of the device

The use of LHSC technique has the advantage of customising the number and size of nodes according to the user requirement. These customisations can be made by changing the amount of ceramic polymer slurry fluid (Fig 1. i), number of holes drilled in the acrylic plate (Fig 1. i), spread area of squeezed fluid (Fig 1.ii) and velocity of separation of two plates (Fig 1.iii). Here we demonstrate that by increasing the number of holes drilled in acrylic plate as mentioned in figure 1.i. a device template with 100 nodes can be obtained (Fig 8.i). The template can further be used to fabricate a high-throughput device with 100 nodes (Fig 8.ii-iii) thus enhancing the resolution of the device.

**Figure 8:**
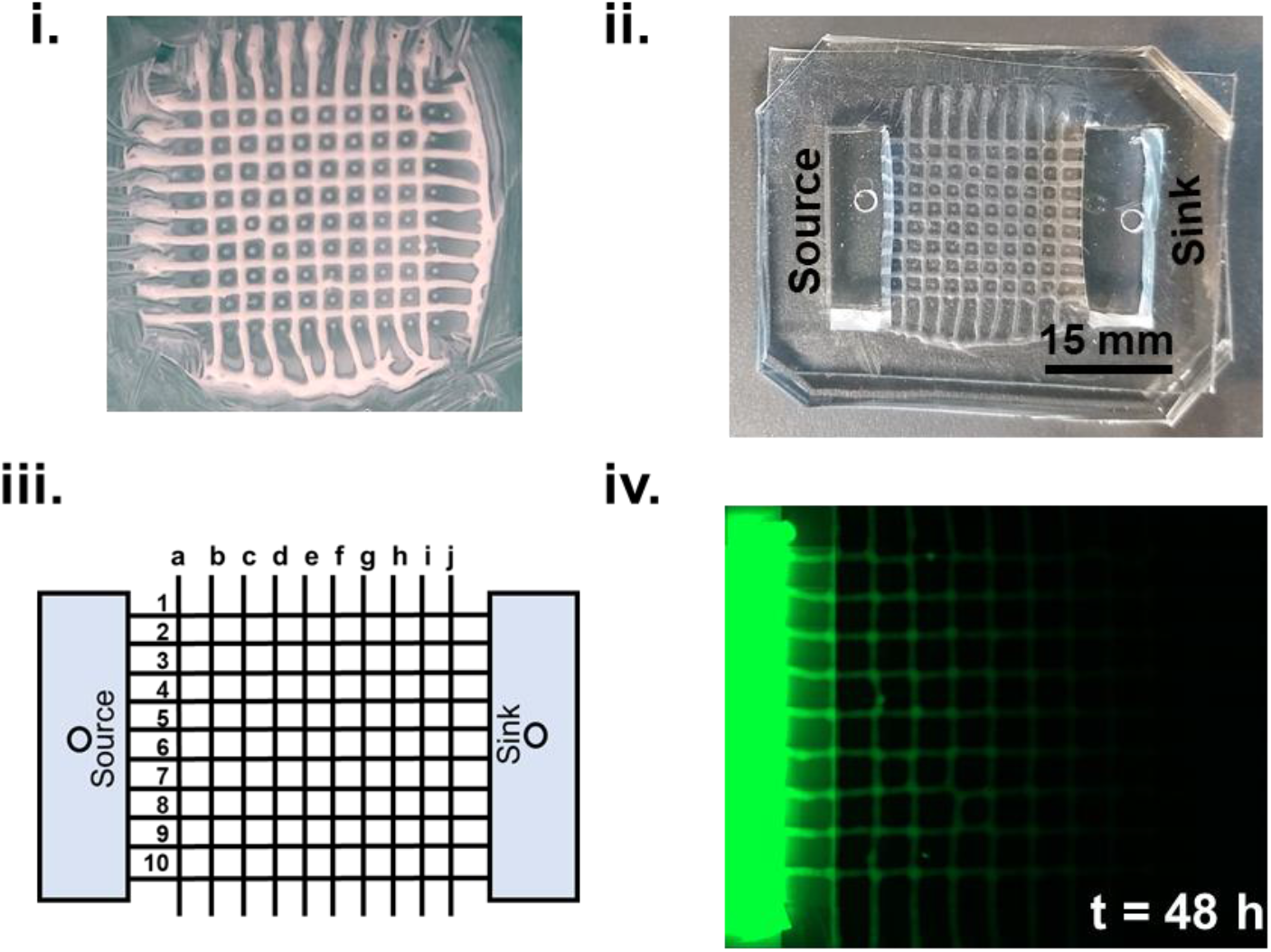
Scalability of the drug testing device: i) Phtograph of template with 100 nodes. ii) Device. iii) Schematic of device with 100 nodes with column channels labelled as a-j from source to sink and row channels as 1-10. iv) FITC gradient in a 9 × 9 device

## 4. Conclusion

In this work, we have demonstrated that the controlled and reproducible shaping of viscous fluids by lifted Hele-Shaw method can be used for fabrication of microfluidic device for drug testing application. To the best of our knowledge this is the first report of the generation of concentration gradient in a flow-less microfluidic device that remains stable for long (> 5 days). As the proposed device is user-friendly, pump- and tubing-less, cost-effective, portable, compatible to the conventional cell culture labs, and can be scaled according to the user’s requirement, it can be potentially adopted and commercialized for wide-scale applications.

## Acknowledgement

We would like to acknowledge the suggestions and technical assistance by Dr. Tanveer ul Islam and Mr. Makrand Rakshe during the fabrication of the template. We want to acknowledge the kind gift of U87-MG cells from Dr. Shilpee Dutt lab, ACTREC, Navi Mumbai and of HeLa cells from the Centre for Cellular & Molecular Biology (CCMB), Hyderabad by Prof. Jyotsna Dhawan. We thank laser confocal imaging facility at IIT Bombay for helping us in procuring the immunostaining images. The work was supported by DST IMPRINT (Department of Science and Technology- Impacting Research Innovation and Technology, Project Number 6722), and WRCB (Wadhwani Research Centre for Bioengineering) IIT Bombay. We also thank DST INSPIRE (Department of Science and Technology- Innovation in Science Pursuit for Inspired Research fellowship) and IRCC (Industrial Research and Consultancy Center) for funding SY and KB’s fellowship respectively.

## Author contribution

AM, PG conceptualized the research and provided overall guidance. KB, SY performed device fabrication and gradient studies. SY performed COMSOL simulations. KB performed all the cellular assays. KB, SY, AM, PG contributed in writing of manuscript.

