## Supplementary material for "Lithography-less, frugal and long-term diffusion-based static gradient generating microfluidic device for high-throughput drug testing": S1-S4

**Keywords**: High throughput drug testing, diffusion-based gradient, microfluidic device

### Supplementary figures-

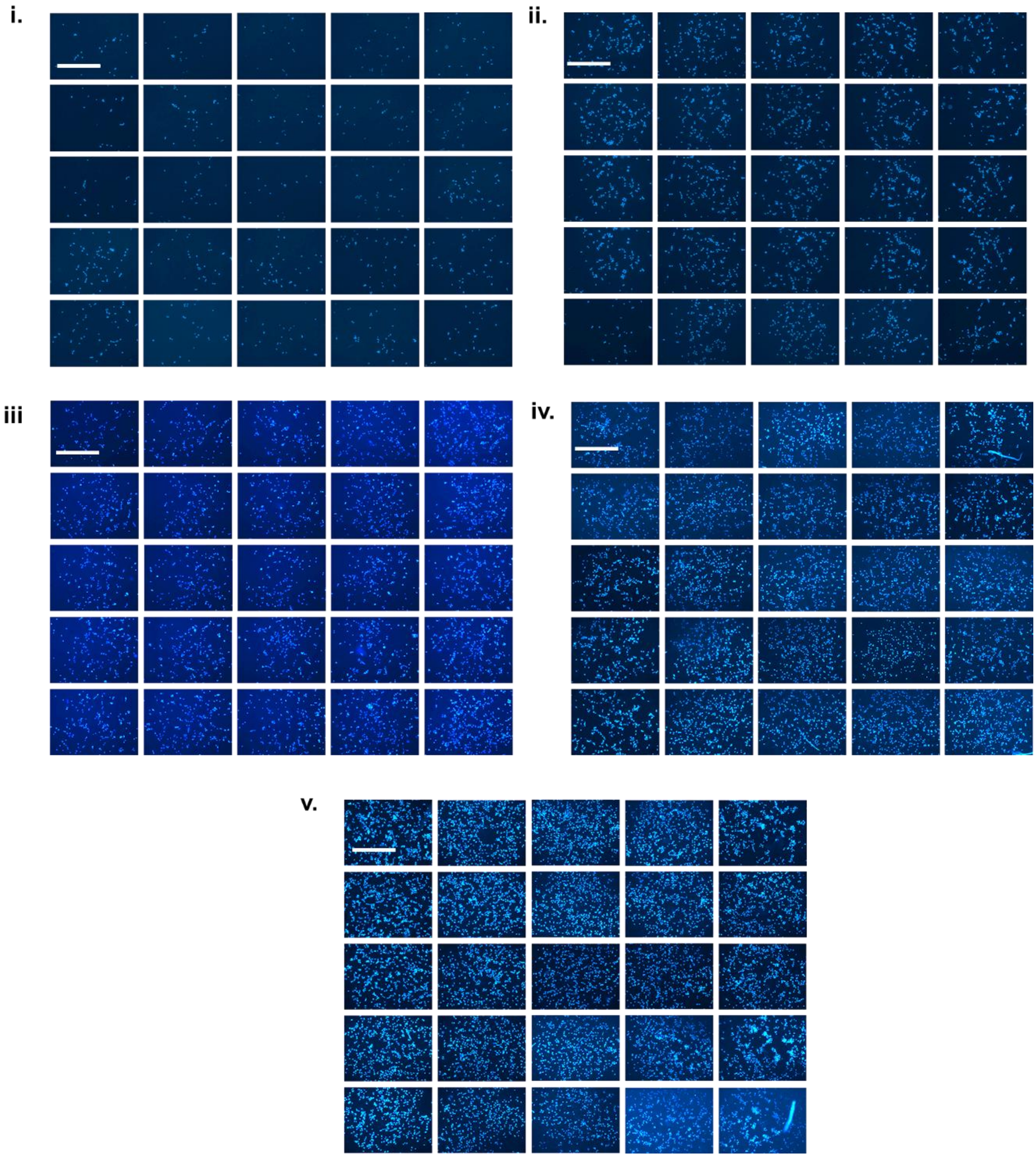


Supplementary figure 1: Equal distribution of cells at the nodes of device represented by hoechst stained cells at 25 nodes of device with seeding density of- i) 0.25 x $10^5$ ii) 0.5 x $10^5$, iii) 1.0 x $10^5$, iv) 2.0 x $10^5$ and v) 4.0 x $10^5$. (Scale bar 400 µm)

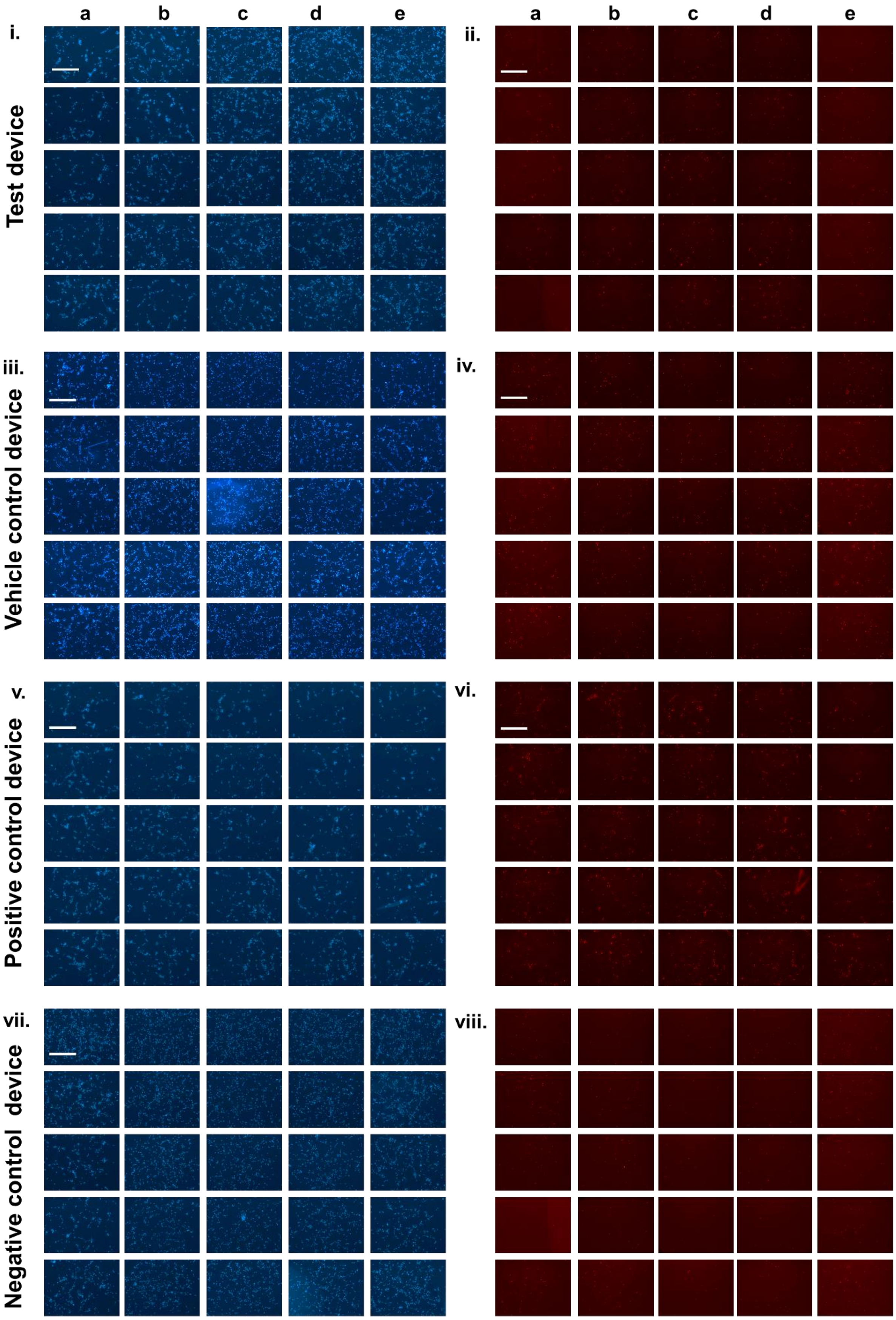
a
b
c
d
e
i.
ii.
Test device
iii.
iv.
Vehicle control device
v.
vi.
Positive control device
vii.
viii.
Negative control device

Supplementary figure 2: Drug testing in device with TMZ. i) Test device with increasing Hoechst stained cells from source to sink (a to e) at 25 nodes ii) Test device with PI stained cells at the corresponding nodes represented in (i), indicating the dead cell population at the 25 nodes of device from source to sink (a-e). iii) DMSO control device with Hoechst staining indicating equal number of cells at all the 25 nodes. iv DMSO control device with PI staining at the corresponding nodes represented in (iii), indicating equal number of dead cells at all the 25 nodes. v) Positive control device with Hoechst stained cells at 25 nodes. vi) Positive control device with PI stained cells at 25 nodes of device. vii) Negative control device with Hoechst stained cells at 25 nodes viii) Negative control device with PI stained cells at the corresponding nodes represented in (vii). (Scale bar 400 µm)

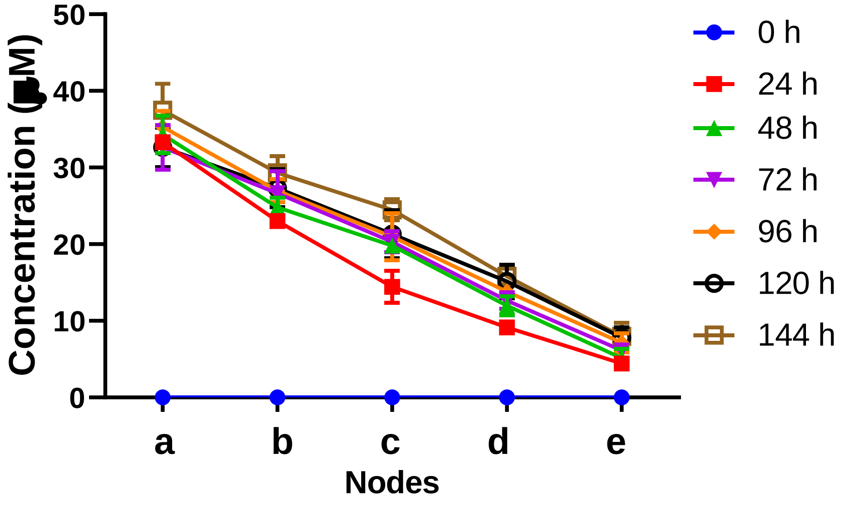


Supplementary figure 3: FITC concentration gradient profile with respect to time (up to 144 h) at different node locations in the microfluidic device.

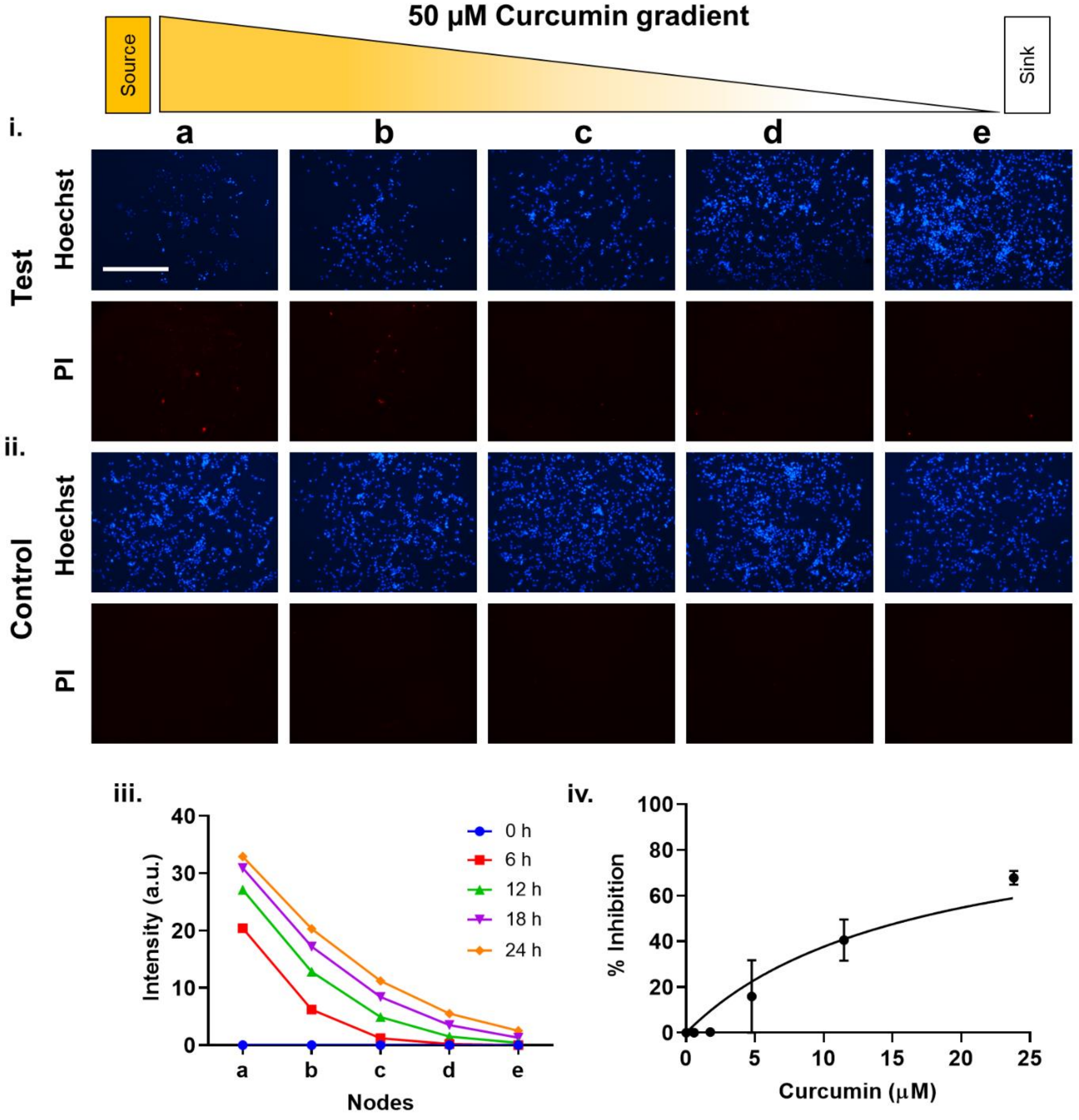


Supplementary figure 4: Estimation of curcumin efficacy using the device: i) Test device: Effect of Curcumin (50 µM) gradient on U87-MG cells was determined using Hoechst and PI staining from source to sink (a-e) in device. ii) Control device: Hoechst and PI staining at 5 nodes (a-e) of the device. iii) Simulation graph of curcumin gradient using COMSOL at 0 h, 24 h, 48 h iv) Percentage inhibition of U87 MG cells at different concentration of curcumin at 5 different nodes from source to sink (a-e) gave $IC_{50}$ as 16.6 µM ± 5.5 µM (Scale bar 400 µm).
